# tRForest: a novel random forest-based algorithm for tRNA-derived fragment target prediction

**DOI:** 10.1101/2021.12.13.472430

**Authors:** Rohan Parikh, Briana Wilson, Laine Marrah, Zhangli Su, Shekhar Saha, Pankaj Kumar, Fenix Huang, Anindya Dutta

**Author notes:** These authors contributed equally (Rohan Parikh, Briana Wilson).

## Abstract

tRNA fragments (tRFs) are small RNAs comparable to the size and function of miRNAs. tRFs are generally Dicer independent, are found associated with Ago, and can repress expression of genes post-transcriptionally. Given that this expands the repertoire of small RNAs capable of post-transcriptional gene expression, it is important to predict tRF targets with confidence. Some attempts have been made to predict tRF targets, but are limited in the scope of tRF classes used in prediction or limited in feature selection. We hypothesized that established miRNA target prediction features applied to tRFs through a random forest machine learning algorithm will immensely improve tRF target prediction. Using this approach, we show significant improvements in tRF target prediction for all classes of tRFs and validate our predictions in two independent cell lines. Finally, Gene Ontology analysis suggests that among the tRFs conserved between mice and humans, the predicted targets are enriched significantly in neuronal function, and we show this specifically for tRF-3009a. These improvements to tRF target prediction further our understanding of tRF function broadly across species and provide avenues for testing novel roles for tRFs in biology. We have created a publicly available website for the targets of tRFs predicted by tRForest.

## Introduction

tRNA fragments (tRFs) are a novel class of small RNAs derived from precursor tRNAs or mature tRNAs (1). tRFs account for nearly 25% of the small RNAs found in the cell and are the second most abundant class of RNAs within the small RNAome. They have been found to play a myriad of roles in normal biology and disease. For example, tRFs have been found to play a role in ribosome biogenesis, respiratory syncytial virus pathogenesis, and breast cancer progression (2–6). More recently, tRFs have been found to augment symbiosis between nitrogen-fixing bacteria and plant hosts via entry of bacterial tRFs into host Ago1 (7).

Following the observation that many tRFs fall within the same size range as microRNAs (miRNAs), we have shown that tRFs can behave as *bona fide* miRNA (8). Until recently, tRF target prediction tools were limited to tools trained on miRNA. This may provide predicted targets that may not be as accurate as algorithms trained on tRFs. Accurate identification of tRF targets will improve our understanding of the role of tRFs in biology and disease. Recognizing this need, tRF specific target prediction tools have become available recently (9–11). Despite the increase in the number of tools over the past couple of years, these tools are lacking in one way or another. For example, tRFtar and tRFTars do not predict targets for tRF-1s, although tRF-1s have been found to affect gene expression in some contexts (12). tRFtarget builds a target algorithm based on RNA intermolecular interactions, but excludes several features that may be important for tRF target prediction, such as target site conservation and AU content (10).

Here we present tRForest, a tRF target prediction algorithm built using the random forest machine learning algorithm. This algorithm predicts targets for all tRFs, including tRF-1s and includes a broad range of features to fully capture tRF-mRNA interaction. Furthermore, unlike other available algorithms, its performance does not rely entirely on just a few features; instead, it uses all available features in ensemble. Finally, since we have employed a machine learning based approach, our predictions can be readily extended to novel tRFs in the future. We used Crosslinking, Ligation, and Sequencing of Hybrids (CLASH) data generated by crosslinking AGO1 with interacting small RNAs and their targets in HEK293 cells as our training and testing dataset (1,13). We rigorously validate the accuracy of tRForest on data from two human cell lines and show that tRForest matches or outperforms currently available tRF and miRNA target prediction tools. We also generated Gene Ontologies (GO) for each tRF’s predicted targets to aid researchers in generating hypotheses about the biological role of each tRF. Existing tRF target prediction tools, tRFtarget and tRFTar have GO analysis capabilities, but are limited in their target prediction features and approach. In addition, our pipeline for producing GO analysis plots is more streamlined, and provides users with publication-ready visualization of the results. tRF predicted targets and GO terms are available to the research community at https://trforest.com/.

## Materials & Methods

### Data and algorithm source

CLASH data with tRFs and their corresponding gene targets were obtained from Kumar et al., which identified all of the tRF-mRNA chimeric reads available in CLASH data from Helwak et al. (1, 13). The tRF sequences and IDs were obtained from tRFdb (http://genome.bioch.virginia.edu/trfdb/) (1). Human gene and transcript sequences, as well as Ensembl and Refseq IDs, were retrieved from Ensembl Biomart (version GRCh37.p13, http://grch37.ensembl.org/biomart/) (14). TargetScan targets were retrieved from TargetScanHuman 7.2 (http://www.targetscan.org/vert_72/) (15). miRDB targets were retrieved from miRDB (http://mirdb.org/) (16). tRFTar predicted targets were obtained from tRFTar (http://www.rnanut.net/tRFTar/) (11). tRFTarget predicted targets were obtained from tRFTarget (http://trftarget.net/) (10). tRFTars predicted targets were obtained from tRFTars (http://trftars.cmuzhenninglab.org:3838/tar/) (9). The random forest classifier used in tRForest was from scikit-learn in Python (https://scikit-learn.org/) (17). RNA-seq data for target validation was obtained from GEO, specifically series GSE99769 (8), series GSE189510, series GSE93717 (18), and series GSE180331.

### RNA-seq library preparation and analysis of RNA-seq data

U87 cells were cultured in MEM supplemented with 1% non-essential amino acids, 1 mM sodium pyruvate, 0.15% sodium bicarbonate and 10% FBS. For RNA mimic transfection, synthetic single-stranded tRF-3009a (5’phos-rArCrCrCrCrArCrUrCrCrUrGrGrUrArCrCrA-3’OH) and non-targeting GL2 control (5’phos-rCrGrUrArCrGrCrGrGrArArUrArCrUrUrCrGrArUrU-3’OH) were transfected at 50 nM final concentration with Lipofectamine 2000 for 48 hours before RNA extraction by ZYMO directzol RNA miniprep kit with DNase I treatment. For mRNA-seq library preparation, 250 or 500 ng total RNA was poly-A selected by NEBNext Poly(A) mRNA Magnetic Isolation Module (NEB #E7490). Library preparation was performed using NEBNext Ultra II Directional RNA Library Prep Kit for Illumina (NEB #7760) according to the manufacturer’s protocol. The resulting libraries were quality checked for concentration and size distribution before pooling for sequencing on Illumina HiSeq by Novogene.

FastQC was used to perform a quality check on the raw RNA-seq data files (https://www.bioinformatics.babraham.ac.uk/projects/fastqc/). Kallisto was then used to quantify the abundances of transcripts using the prebuilt ENSEMBL index available in the kallisto manual (19). DESeq2 was then utilized in R to perform differential expression analysis with an expression cutoff of 10 normalized counts (20). Finally, BioMart was used to convert Ensembl transcript IDs to gene names (14).

### miRNA and tRF prediction algorithms

TargetScan and miRDB were used to find the predicted gene targets for a given tRF or miRNA, such as tRF-3009a or miR-941, with the following search parameters: for TargetScan and miRDB, the default parameters were used, with 7mer-m8 seed pairing for TargetScan and the full tRF sequence for miRDB. For each algorithm, predicted targets were separated from non-targets. The log2 fold changes of targets and non-targets were used to generate cumulative distribution function (CDF) plots. This was done for available tRF prediction algorithms as well, such as tRFTar, tRFTarget, tRFTars, and tRForest, with the following search parameters: for tRFTar, 3’ UTR target element type with no restrictions on the expression levels of the tRF or co-expression of tRF-gene pairs; for tRFTarget, 3’ UTR binding region and default free energy (≤ −10 kcal/mol) and maximum complementary length (≥ 8 nts); for tRFTars, the SVM-GA model with high confidence; for tRForest, targets with a prediction score of ≥ 0.8. Only 3’ UTR targets were compared because tRForest is only trained on and predicts targets in the 3’ UTR.

### Generation of negative sites

The CLASH data only provided positive ground-truth data to train the random forest with; we classified the targets found in the CLASH data as ground-truth because they were experimentally validated. In order for the training to be most effective, realistic negative sites had to be generated corresponding to each positive site. This was done similarly to TarPmiR (21). For each positive ground-truth tRF-mRNA duplex in the CLASH data, a set of candidate negative sites was first generated by running a sliding window of the same length as the binding region on the 3’ UTR of the mRNA upstream and downstream of the binding region. Then, the CG dinucleotide frequency in the binding region was identified; if the dinucleotide frequency was the same as the corresponding positive binding site, the frequency of the G nucleotide was then identified. If the frequency of the G nucleotide was also similar between the candidate negative site and corresponding positive site, it passed the first filter. After filtering the original list with the CG dinucleotide and G nucleotide frequencies, the binding energy between the tRF and each of the remaining candidate negative sites was calculated, and the candidate site with the lowest energy was selected as the negative corresponding site. These filters were used when generating the negative site to allow the negative site to be sufficiently similar to the positive site, so that the random forest would distinguish them using the calculated features, and not exclusively binding energy or because of the frequency of individual nucleotides (22).

### Feature calculation

Thirteen features were calculated for each tRF-mRNA ground-truth duplex in the CLASH dataset. These included: (i) binding energy; (ii) seed match; (iii) AU content; (iv) number of paired positions; (v) binding region length; (vi) the length of the longest consecutive pairing; (vii) the position of the longest consecutive pairing; (viii) the number of 3’ end pairs; (ix) seed-3’ end pair difference; (x) binding region conservation with phyloP scores; (xi) flanking region conservation with phyloP scores; (xii) binding region conservation with phastCons scores; and (xiii) flanking region conservation with phastCons scores. These features were selected due to their experimental importance as exemplified by TargetScan, as well as their successful use in TarPmiR, a random forests-based target prediction algorithm for microRNA (19). In addition, we previously established that tRFs can downregulate mRNA targets in a miRNA-like mechanism, using seed sequences (23). Therefore, we used modified miRNA seed binding rules as a feature in tRForest. The only modification was that we allowed binding at the first nucleotide of the tRF. The detailed definitions and information on the calculation of each of these features is described in supplementary file 1.

### Random forest training and testing

Scikit-learn’s RandomForestClassifier() function was used to train and get testing metrics for tRForest (17). tRForest was trained in two ways to obtain metrics of its performance: 10-fold cross validation (in which the training data was split into 10 equal blocks of data, with 9 blocks of data being used for training and 1 block for testing, iterating until each block had been used as a testing set) and with a classic 67:33 train-test split. From this, the following performance metrics were obtained: accuracy, positive predictive value (PPV), sensitivity, F1 scores, and a receiver operating characteristic (ROC) curve with the area under the curve (AUC). Hyperparameter tuning was performed for algorithm optimization. We tested several values for the number of trees in the random forest, the number of features to consider when looking for the best split, the maximum depth of the tree, the minimum number of samples required to split an internal node, the minimum number of samples required to be at a leaf node, and whether bootstrapping was conducted. However, after varying the hyperparameters, a majority of them were kept at their default setting, barring two. The number of trees in the random forest was doubled to 200 trees, and bootstrapping was disabled.

### Algorithm robustness

Several tests were performed to test the robustness of the trained algorithm. First, the Pearson correlations between the features were examined for the feature profiles of tRF-3009a, as the random forests algorithm is most effective when its features have very low internal correlation (24). Pearson correlations were chosen because the correlation value represents the linear relationship between continuous variables. After this, the algorithm was tested with various subsets of features in order to determine whether the choice to classify a gene as a target for a particular tRF hinged on just one highly important feature or a small subset of important features, or whether it was using an ensemble of all of the features. This was done by removing each feature, one at a time, and then by removing the 7 most important and least important features, where the importance of the features was ranked through recursive feature elimination (RFE() function from Scikit-learn).

Finally, an accuracy-efficiency analysis was performed, in which the effect of dropping the most time-intensive features on the performance of the algorithm was determined, in order to create the most efficient algorithm possible. This was done by ranking the features in order of time required to calculate them, removing these features one at a time from the most time-intensive to least time-intensive, and then determining the performance metrics of the algorithm. Based on this analysis, secondary structure accessibility, phastCons stem conservation score, and phastCons flanking regions conservation score were removed (phyloP conservation scores were kept), resulting in a total of 11 features in the final iteration of tRForest.

### Algorithm validation

To validate the algorithm, tRForest was tested on two independent RNA-sequencing datasets following experimental perturbation of the abundance of the tRFs. This was done by creating the feature profile for the perturbed tRF, and then determining the predicted targets. After this, the genes from the RNA sequencing dataset were separated into targets and non-targets, and CDF plots were generated comparing the log2 fold change of targets compared to non-targets to determine whether the targets were repressed when the tRF was increased.

These CDF plots were compared to those generated from the predicted targets of tRFTar, tRFTarget, and tRFTars, three existing tRF prediction algorithms, as well as miRDB and TargetScan, two existing miRNA prediction algorithms. Effect sizes (X-axis displacement between the two CDF plots: Log2 fold change of targets at 0.5 fraction of genes – Log2 fold change of non-targets at 0.5 fraction of genes) and the two-sided p-values from a Kolmogorov-Smirnov (KS) test were also calculated.

### tRF gene target database

After the algorithm had been validated, tRForest was used to predict gene targets for each of the tRFs in tRFdb across seven species, excluding the tRFs from *R. sphaeroides* because it is the only species in the database that does not undergo eukaryotic Ago-mediated repression (1, 25). This was done by first generating a list of all transcripts in each genome (human: version GRCh37.p13/hg19; mouse: mm9; *D. melanogaster*: dm3; *C. elegans*: ce6; *S. pombe*: schiPomb1; Xenopus: xenTro3; zebrafish: Zv9). For each tRF, two passes were first conducted: a seed match pass and a binding energy pass. In miRNA, the first nucleotide is generally unavailable for binding due to the architecture of the Argonaute protein; however, we considered the first position of the tRF to determine whether it was a factor in target prediction (26). Thus, for the first pass, a list of transcripts with 7mer-m1 seed matches (canonical pairing in nucleotides 1-7 of the tRF) to the tRF was generated. For the second pass, the binding energy of a tRF to each transcript was calculated, and a list of transcripts with a more stable binding energy than 60% of the binding energy from a perfect tRF-mRNA duplex was generated. For the transcripts in the intersection of the two passes, the remaining 9 features were calculated and the feature profiles for the tRF-mRNA duplexes were passed to tRForest, which determined the final predicted targets. For *S. pombe*, the fission yeast species, conservation scores were not calculated. RNAhybrid was also used to create interaction illustrations of the tRF-mRNA duplexes (27). These targets were placed in a database, which can be queried through different criteria including tRF type, tRF ID, gene name, and transcript ID at https://trforest.com. The tool returns the Ensembl transcript ID, gene name, binding energy, binding location on the 3’ UTR, and an interaction illustration for each target, along with an option to return the chromosomal coordinates of the 3’ UTR of each target.

### Gene ontology analysis

Gene ontology analysis was implemented using the clusterProfiler package https://bioconductor.org/packages/release/bioc/html/clusterProfiler.html (28). First, the list of tRForest predicted genes was converted to Entrez IDs using BioMart (14). Then, the gene list was processed for each of the three different ontology types (Biological Process, Molecular Function, and Cellular Component) using the enrichGO() function, which yielded pathways ranked based on number of genes relating to the pathway and the adjusted p-value. These enrichment outputs were then used to create two different plots for each ontology type. First, a dotplot was created using the dotplot() function, which showed the ten pathways with the highest gene ratios and displayed data for the gene ratio, gene count, and adjusted p-value. Secondly, a network plot was created using the cnetplot() function, which showed the connections between various genes and the highest-ranking pathways. These two plots for each ontology type were then combined into a final figure containing six plots for each tRF and each species. For some tRFs, though, one or more ontologies would not have enough genes to generate an output, in which case there would be fewer plots for that tRF.

For two species (*Xenopus tropicalis* and *Schizosaccharomyces pombe*), the annotations were not already part of the organism database (OrgDb) built in as part of clusterProfiler, so extra steps were taken to build the databases. For *X. tropicalis*, species annotations were available within the AnnotationHub resource https://bioconductor.org/packages/release/bioc/html/AnnotationHub.html, so those annotations were downloaded within R and used as an OrgDb. For *S. pombe*, annotations were created using the makeOrgPackageFromNCBI() function in the AnnotationForge package https://bioconductor.org/packages/release/bioc/html/AnnotationForge.html, which downloaded annotations directly from NCBI and stored them locally in order to build an OrgDb. From here, these two OrgDbs were used just as the built-in ones were when performing the GO analysis.

### Code with documentation

The code for tRForest is available at https://github.com/tRForest-team/tRForest.git.

## Results

### tRForest metrics & algorithm robustness

Figure 1 shows a visual representation of the structure of a random forest and how it is used for classification of small RNA targets and non-targets. Figure 2 shows a comprehensive workflow describing the development of the tRForest algorithm. We obtained ground-truth targets from an AGO1 cross-linking, ligation, and sequencing of hybrids (CLASH) dataset (1) and used them to train tRForest using the calculated features described in Methods. After several optimization steps described below, the final algorithm was obtained.

**Figure 1.**
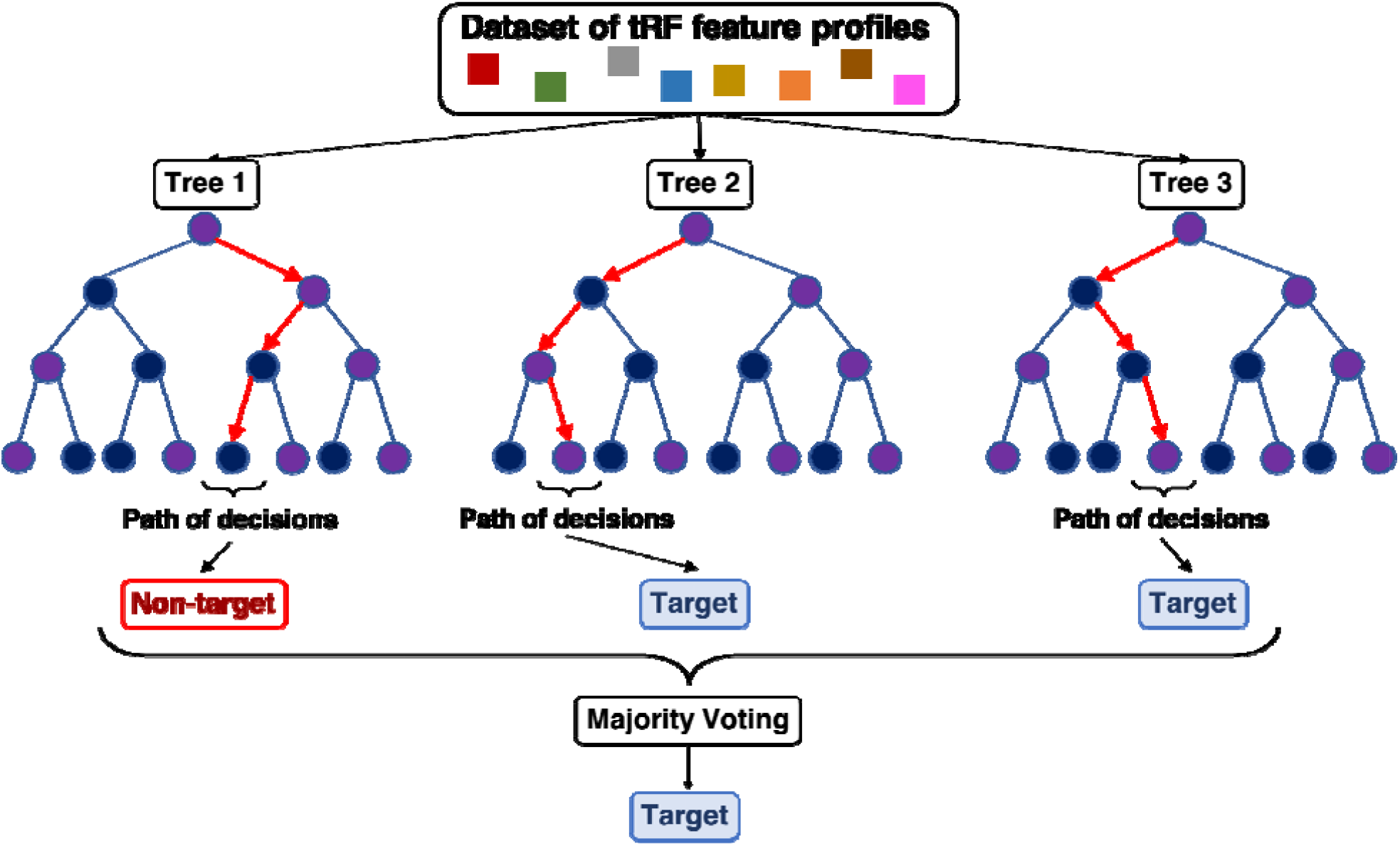
An example structure of a random forest algorithm, including how the forest is used for classification. Feature profiles (squares) represent each tRF-gene pair’s unique set of feature calculations for all 11 features. These feature profiles are fed through independent trees which each make a series of binary decisions at each node (circle). To make this decision at each node, the tree identifies the best feature for optimal splitting of the node from a subset of random features from the feature profile (29). Over the course of several such splits, the tree gains a better understanding of the tRF-gene pair. This series of decisions results in the tree classifying the gene as a target or non-target, and the majority vote of all trees in the algorithm determines the overall classification.

**Figure 2.**
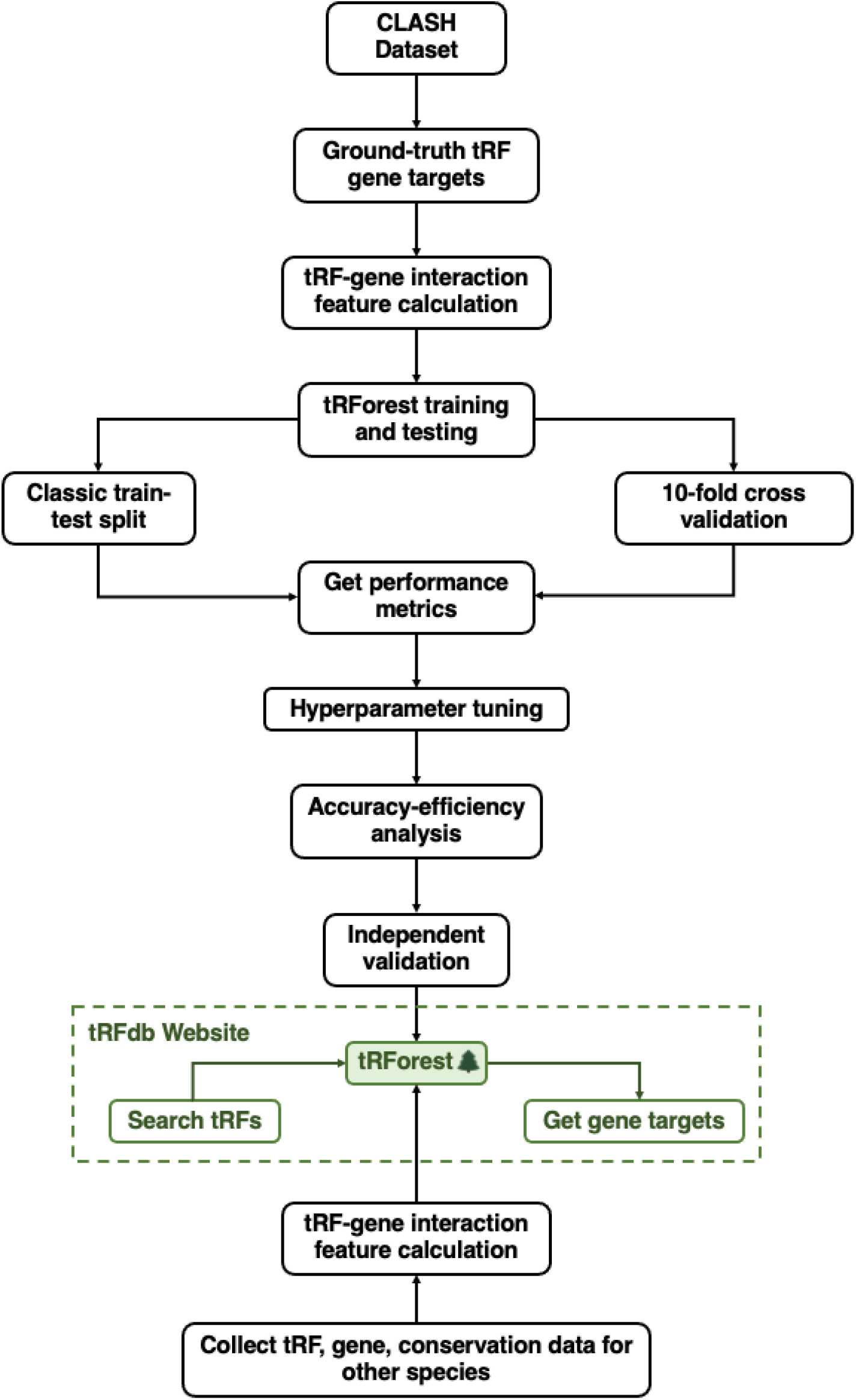
Comprehensive workflow from initial dataset to validated algorithm applied to tRFs in seven species.

To decide on our final optimized algorithm, we first determined the sensitivity and positive predictive value (PPV) of tRForest. Sensitivity (calculated as true positive / (true positive + false negative)) is the ability to detect positive targets and PPV (calculated as true positives / (true positives + false positives) is the ability to detect true positive targets among all predicted positive targets. Together, these metrics show the algorithm is both sensitive and specific. The algorithm performs best with 7mer-m1 or 8mer seed matching (Table 1; see Figure 3 for seed matching schematic). It is possible that 7mer-m1 matching outperforms other 7mer seed matching criteria because tRFs, unlike miRNAs, can actually pair with mRNA in the first position. However, it may also be due to the fact that adenine is the first nucleotide in approximately 50% of tRFs, making the seed match identical to the 7mer-A1 criteria in 50% of cases (supplemental figure 1). In tRForest, 7mer-m1 seed matching was used because it provided a larger number of high-confidence targets than 8mer seed matches. Sensitivity and PPV were 0.9616 and 0.9620 respectively when 7mer-m1 seed matches were used and when considering all features (Table 2).

**Table 1.**
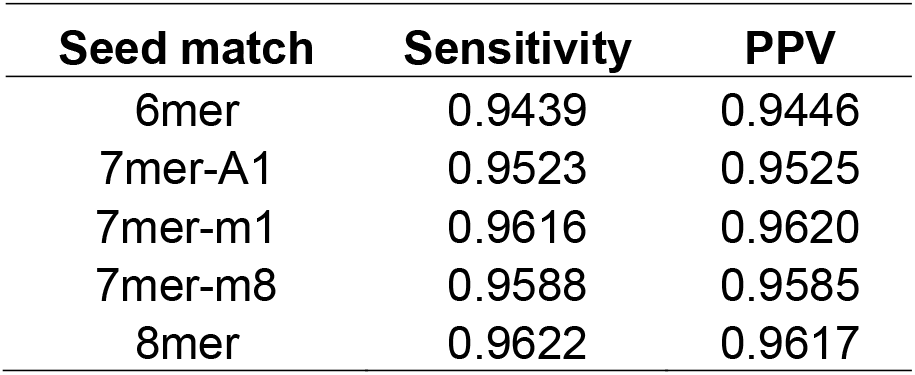
Sensitivity and PPV values for tRForest with different seed match criteria (see Figure 3). 7mer-m1 and 8mer seed matches outperform other criteria.

**Table 2.** Run time comparisons, sensitivity and PPV values for tRForest evaluated on various subsets of features. The run times are ranked from 0 (slowest) to 13 (fastest). The clustering of the sensitivity and PPV values after removal of various features suggests that no individual or small subset of features contributes disproportionately to the classification. RFE: Recursive Feature Elimination. Last row: tRForest performs significantly better when labels properly indicate targets and non-targets compared to randomly shuffling the labels.

| Rank by time to calculate | Feature subset | Sensitivity | PPV |
| --- | --- | --- | --- |
| 0 | All features | 0.9616 | 0.9620 |
| 1 | Removed phastCons flanking score | 0.9596 | 0.9597 |
| 2 | Removed phastCons stem score | 0.9603 | 0.9605 |
| 3 | Removed phyloP flanking score | 0.9596 | 0.9599 |
| 4 | Removed phyloP stem score | 0.9594 | 0.9597 |
| 5 | Removed binding energy | 0.9583 | 0.9587 |
| 6 | Removed position of longest consecutive pairing | 0.9606 | 0.9609 |
| 7 | Removed length of longest consecutive pairing | 0.9605 | 0.9610 |
| 8 | Removed AU content | 0.9604 | 0.9608 |
| 9 | Removed number of paired positions | 0.9616 | 0.9620 |
| 10 | Removed seed | 0.9602 | 0.9607 |
| 11 | Removed binding region length | 0.9570 | 0.9577 |
| 12 | Removed number of 3' end pairs | 0.9602 | 0.9605 |
| 13 | Removed difference between seed and 3' end pairs | 0.9613 | 0.9616 |
| N/A | All RFE ranked > 1 (RFE = 7) | 0.9458 | 0.9453 |
| N/A | All RFE ranked = 1 (RFE = 7) | 0.9423 | 0.9422 |
| N/A | Randomly shuffled | 0.5053 | 0.5054 |

**Figure 3.**
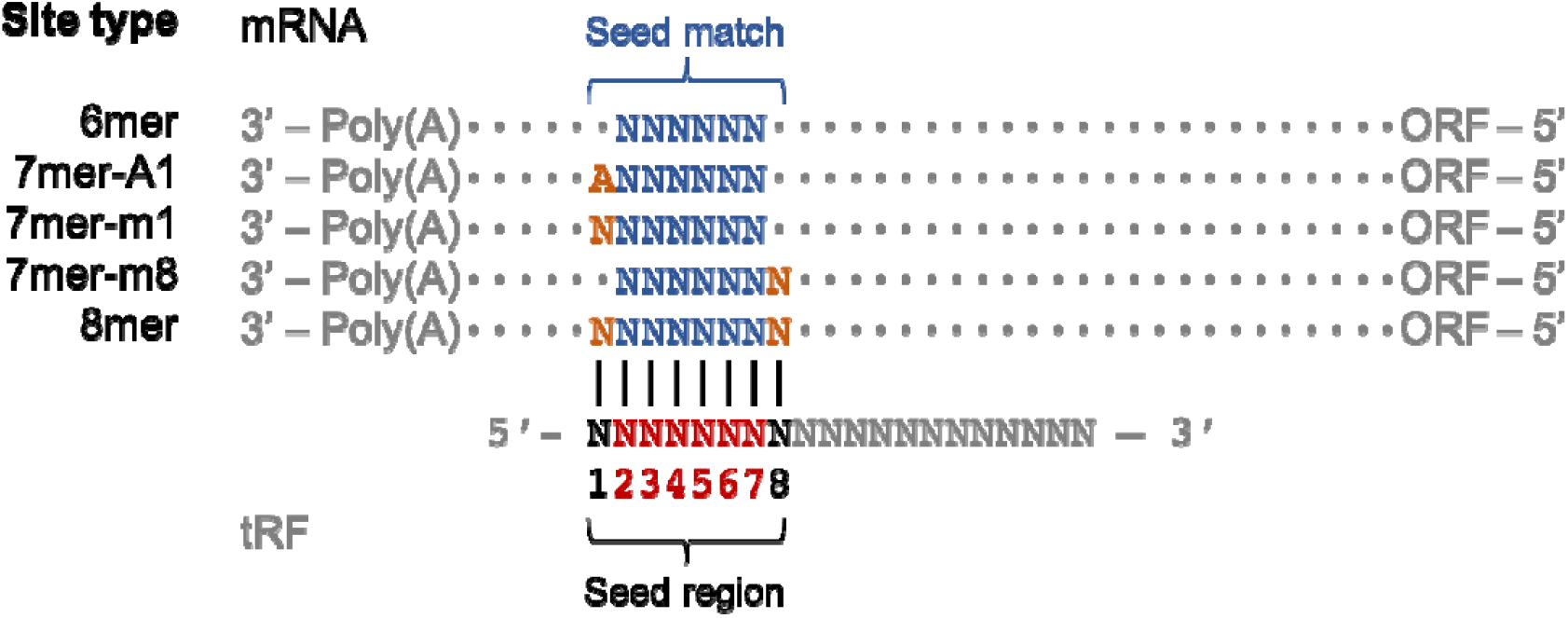
Site type breakdown for seed-matching. The 6mer seed comprises nucleotides 2-7 on the tRF, with additional matching of the target at nucleotides 1 and/or 8 on the tRF based on the site type.

Random forest algorithms work optimally when there is little correlation between features. Therefore, we determined the correlation between each feature in tRForest using tRF-3009a as an example (Figure 4). The magnitude of the correlation was close to zero for a majority of the pairs. There were two groups of pairs with high correlation. The first group consisted of pairs in which information about one feature was used in the calculation of the other feature. For example, there was a strong correlation (r = 0.590) between the presence of a seed in the binding region and the difference in the number of paired positions between the seed region and 3’ region of the tRF. In addition, there was a strong correlation (r = 0.677) between the number of paired positions and the length of the longest consecutive sequence of pairs in the binding region. The features in both of these pairs were kept in the algorithm because although there was a moderate correlation, they provided distinct pieces of information about the tRF-mRNA duplex. The second group of correlated pairs consisted of two sets of conservation scores of the binding region and the flanking region, phastCons scores and phyloP scores. These scores differ in that phastCons scores give the probability that a nucleotide belongs to a conserved element, while phyloP scores give the logarithm of the p-value under a null hypothesis of neutral evolution. From this analysis, we decided to remove the phastCons scores and keep the phyloP scores, because phyloP scores provide information on both slower-than-neutral and faster-than-neutral evolution.

**Figure 4.**
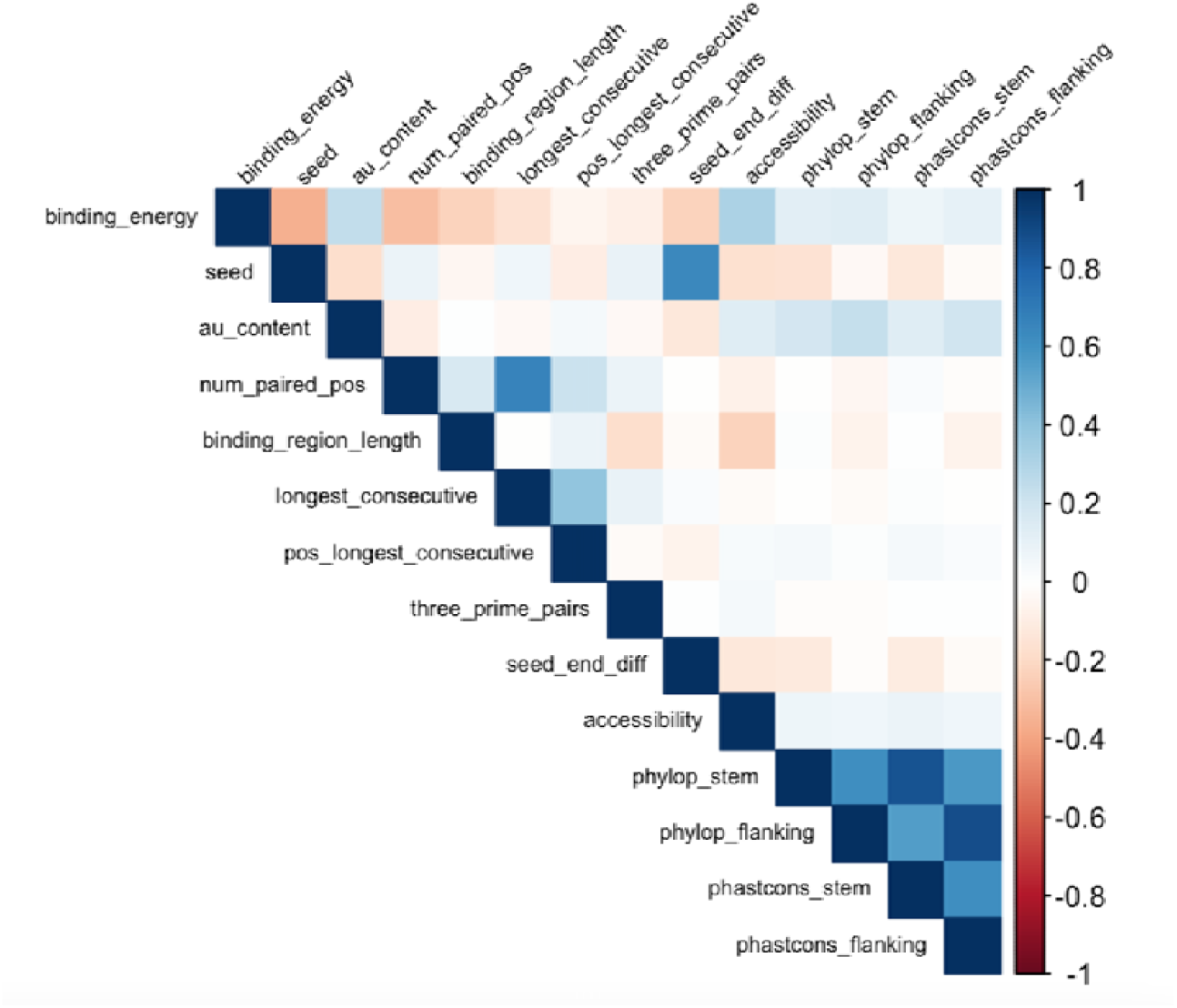
Heatmap of Pearson correlations between pairs of features for tRF-3009a. Only the two different methods of computing evolutionary conservation scores are highly correlated, with low-to-moderate correlation between all other pairs.

In order to reduce the time to run tRForest, we performed an accuracy-efficiency analysis in which the accuracy of the algorithm was measured as time-intensive features were removed (Figure 5). The ranking of the features according to time to calculate is in Table 2. We found that the accuracy remained above 95% after the two most time-intensive features, the phastCon conservation scores (seed and flanking), were removed. Thus, both the correlation and accuracy-efficiency analyses support the removal of the phastCons scores, allowing tRForest to have low-to-moderate correlation between features, maintain an accuracy above 95%, and achieve PPV and sensitivity values of 0.96.

**Figure 5.**
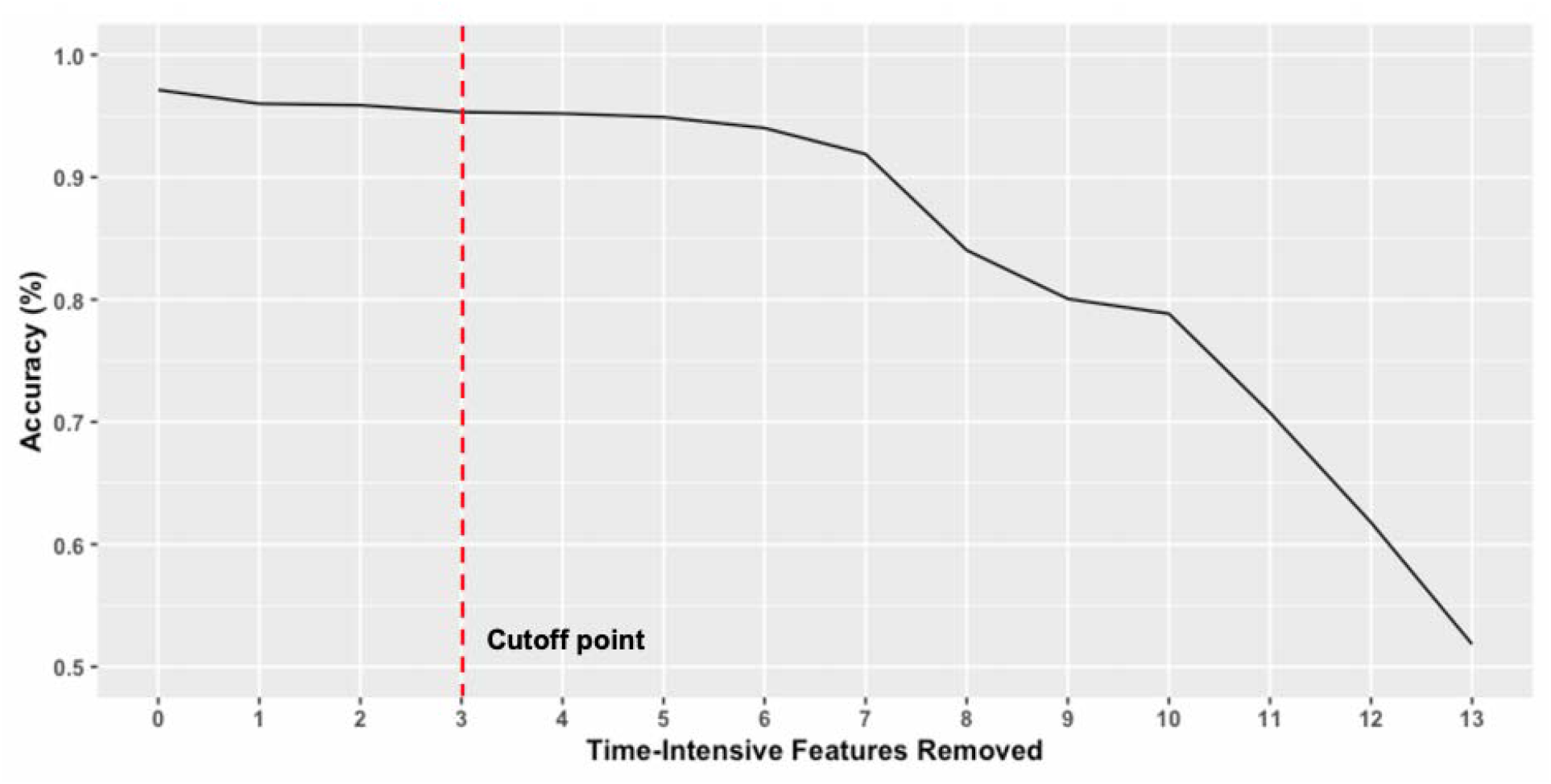
Accuracy-efficiency analysis of tRForest. tRForest continues to perform well as the most time-intensive features are removed, allowing it to become more efficient while maintaining accuracy above 95%.

To ensure robustness of the algorithm, we first removed individual features from the training set. Even when half of the most important or least important features are removed, the algorithm performs effectively (PPV = ~0.944, sensitivity = ~0.944), indicating that no feature or set of features is much more important than any other features (Table 2). Since these features and feature subsets were able to sufficiently train the algorithm, additional features did not need to be added. Furthermore, many of these features are easily calculated, increasing the efficiency of the algorithm. To ensure the algorithm was not merely classifying all targets as positive targets and to ensure the negative sites were properly generated, the labels on the positive and negative sites were randomly shuffled. When the algorithm was trained this way, the PPV and sensitivity dropped to ~0.5, indicating the algorithm was simply guessing (Table 2). Therefore, tRForest performs significantly better than random guessing.

Finally, a receiver operating characteristic (ROC) curve was also generated from the 10-fold cross validation training of the optimized tRForest (after the phastCons conservation features were removed). The ROC curve for tRForest is very close to an ideal classifier and has an area under the curve (AUC) of 0.99, in which the ideal value is 1 (Figure 6).

**Figure 6.**
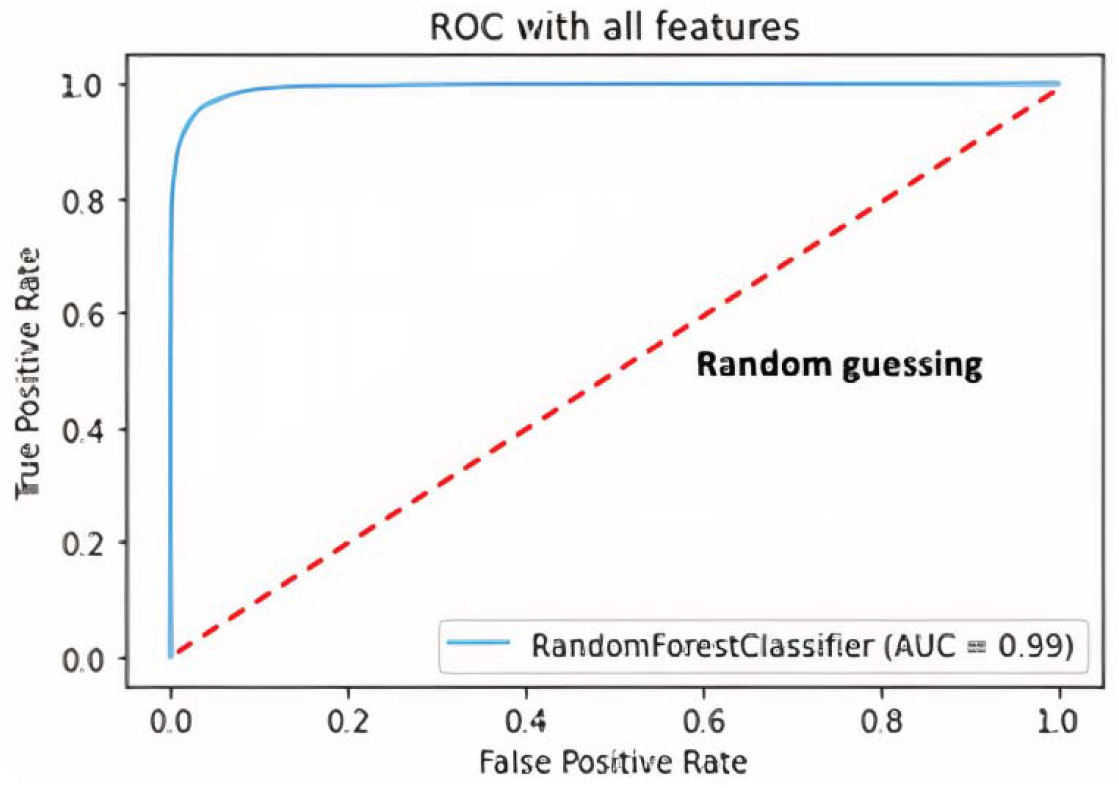
Receiver operating characteristic curve. tRForest performs nearly ideally compared to random guessing during training and testing.

### Independent validation & comparison to existing miRNA and tRF algorithms

Next, we wanted to determine how well tRForest performs on new data derived from tRF overexpression. We used two independent datasets with two separate experimental conditions in two different cell lines. First, we analyzed data in which tRNA-LeuTAA was overexpressed in HEK293T cells, which led to increased tRF-3009a expression (8). We observed that tRF-3009a targets predicted by tRForest were significantly repressed compared to non-targets by cumulative distribution function (CDF) plot (Figure 7; KS test p value = 3.65E-04). The next dataset we analyzed was from overexpression of a tRF-3009a mimic in U87 cells. Again, we observed that tRF-3009a targets predicted by tRForest were significantly repressed compared to non-targets by CDF plot (Figure 8; KS test p value = 2.64E-05).

**Figure 7.**
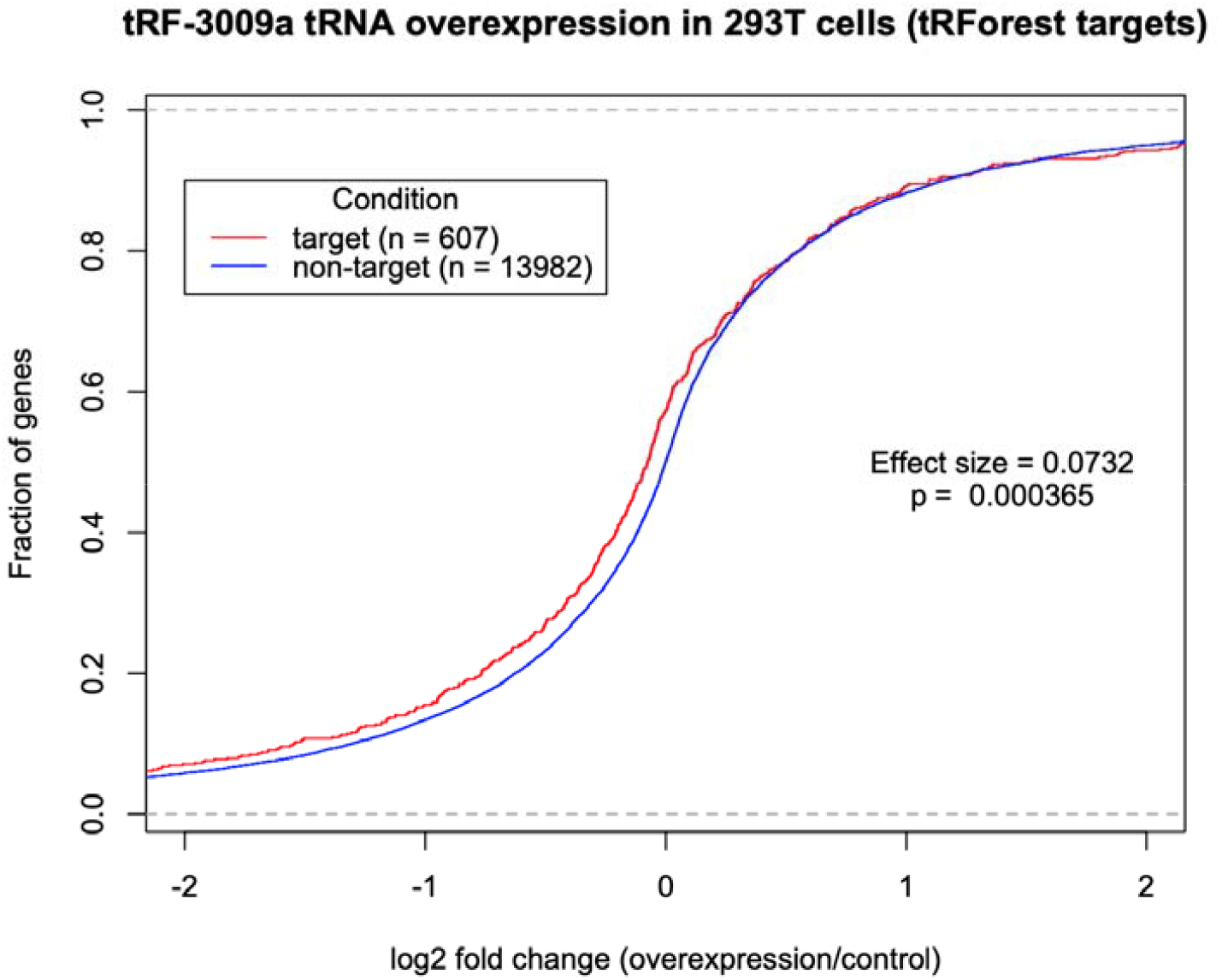
tRForest distinguishes targets from non-targets of tRF-3009 in RNA-sequencing data collected after parental tRNA (chr6.trna83-LeuTAA) overexpression in HEK293T cells. This is known to increase the level of tRF-3009a (8).

**Figure 8.**
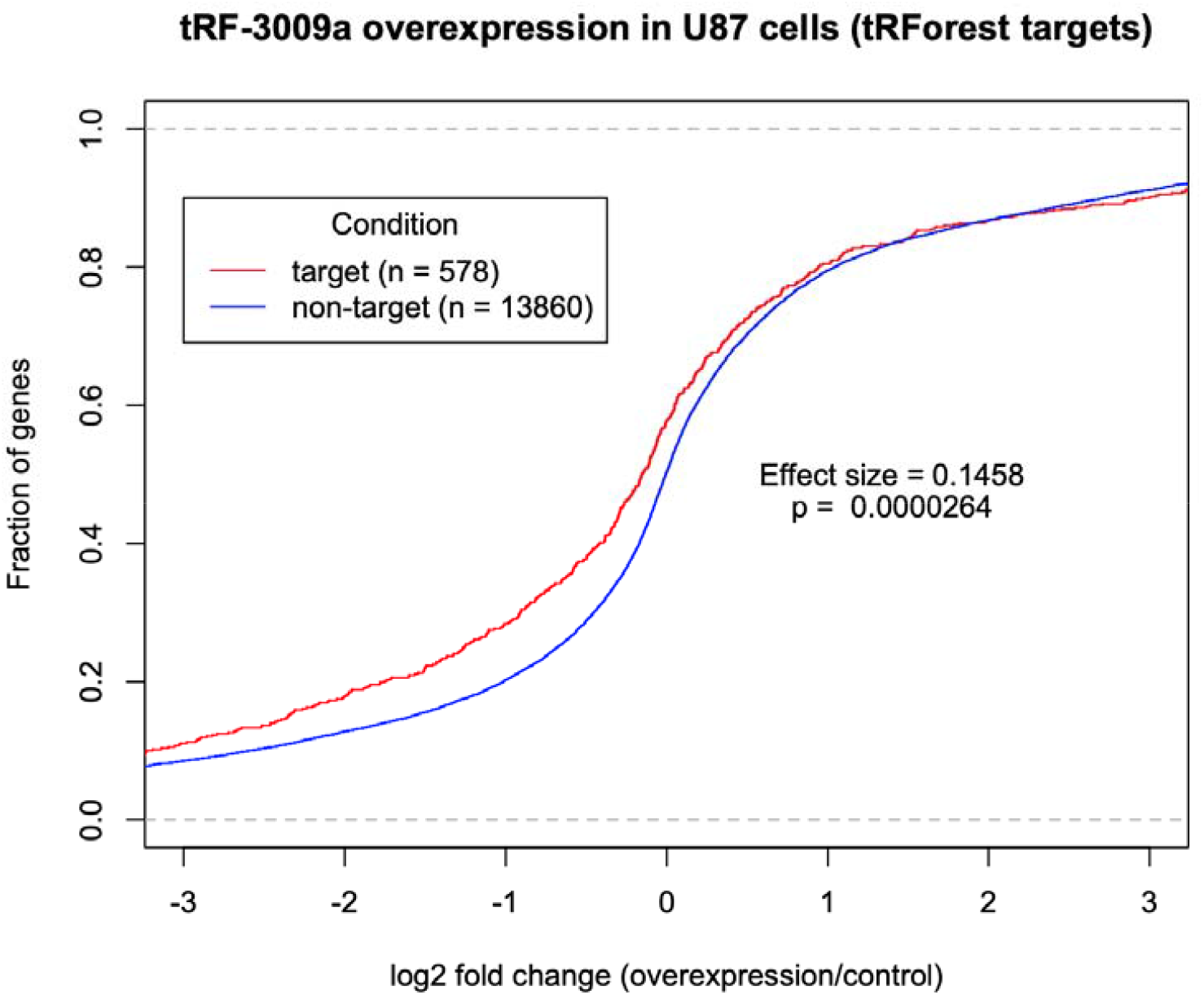
tRForest distinguishes targets from non-targets of tRF-3009 in independent RNA-sequencing data collected from upon tRF-3009 mimic single-stranded overexpression transfection in U87 cells.

CDF plots for targets predicted by other miRNA and tRF algorithms are in supplemental figures 2 and 3. For each cell type, the predicted targets from miRDB, targetScan, tRFTar, tRFTarget, tRFTars, and tRForest were compared to non-targets. We established an effect size statistic to compare the difference between targets and non-targets for the different algorithms (see methods). A negative effect size indicates target repression. Table 3 shows effect sizes and p-values from each validation experiment. tRForest outperforms miRNA target prediction algorithms overwhelmingly with much larger effect sizes and by showing significance with p < tRForest also outperforms tRFTarget and tRFTars with respect to effect size by nearly a factor of two in the HEK293T cell experiment, and by a factor of two to four in the U87 cell experiment with p < 0.001. tRForest matches the performance of tRFTar with similar effect sizes, but presents a greater number of targets with much greater significance (p < 0.001). The range of effect sizes and number of predicted targets from tRForest are comparable to miRDB and targetScan evaluated on miRNA mimic overexpression data, indicating that with the appropriate target prediction tool, tRFs are just as functional as miRNAs, at least in the case of tRF-3009a. Overall, tRForest either closely matches or greatly outperforms all existing tRF target prediction algorithms along with several miRNA target prediction algorithms, and matches the performance of established miRNA algorithms on miRNA overexpression datasets.

**Table 3.**
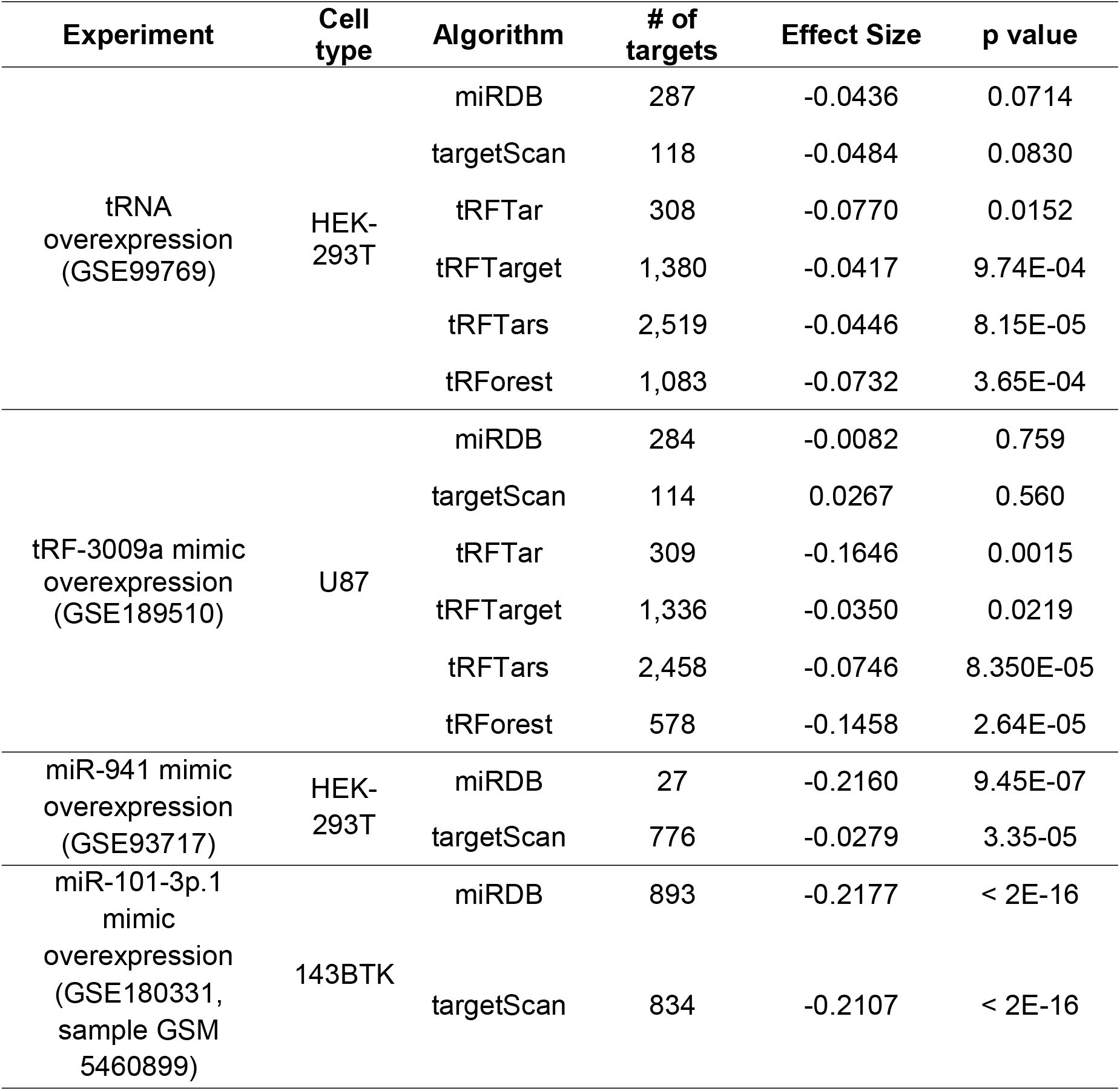
tRForest outperforms several miRNA target prediction algorithms in its ability to predict repression of target genes upon tRF induction in two different cells. tRForest matches or outperforms all existing tRF target prediction algorithms in its ability to predict repression of target genes upon tRF induction in two different cells. Its performance is also comparable to miRDB and TargetScan evaluated on miRNA mimic overexpression data.

### Gene ontology analysis

Following target prediction and validation, gene ontology analysis was performed for each tRF in order to provide an overview of pathways potentially affected by a given tRF. Strikingly, we identified neural and axonogenesis related GO terms in the tRF targets as top pathways enriched across human, mouse, zebrafish, and drosophila (Figure 9A). To determine if these or other GO terms are conserved across these species, we first identified conserved tRF sequences across these species. The highest number of conserved tRFs were found between human and mouse, with 54 tRFs conserved at the sequence level (Figure 9B). Human, mouse, and drosophila have 3 conserved tRFs, and human, mouse, and C. elegans have only one conserved tRF (Figure 9B). Since humans and mice had the highest number of conserved tRFs, we sought to determine which GO terms are conserved for the conserved tRFs in these species. We identified 409 conserved GO terms between mice and humans (Figure 9C). Of particular interest, the top conserved GO terms between mice and humans involve brain related processes, indicating that tRFs may have a conserved role in regulating genes in the nervous system. Figure 9E shows an example output for the human tRF-3009a Biological Process ontology. This includes both the dotplot and the gene concept network plot as generated by clusterProfiler (28).

**Figure 9.**
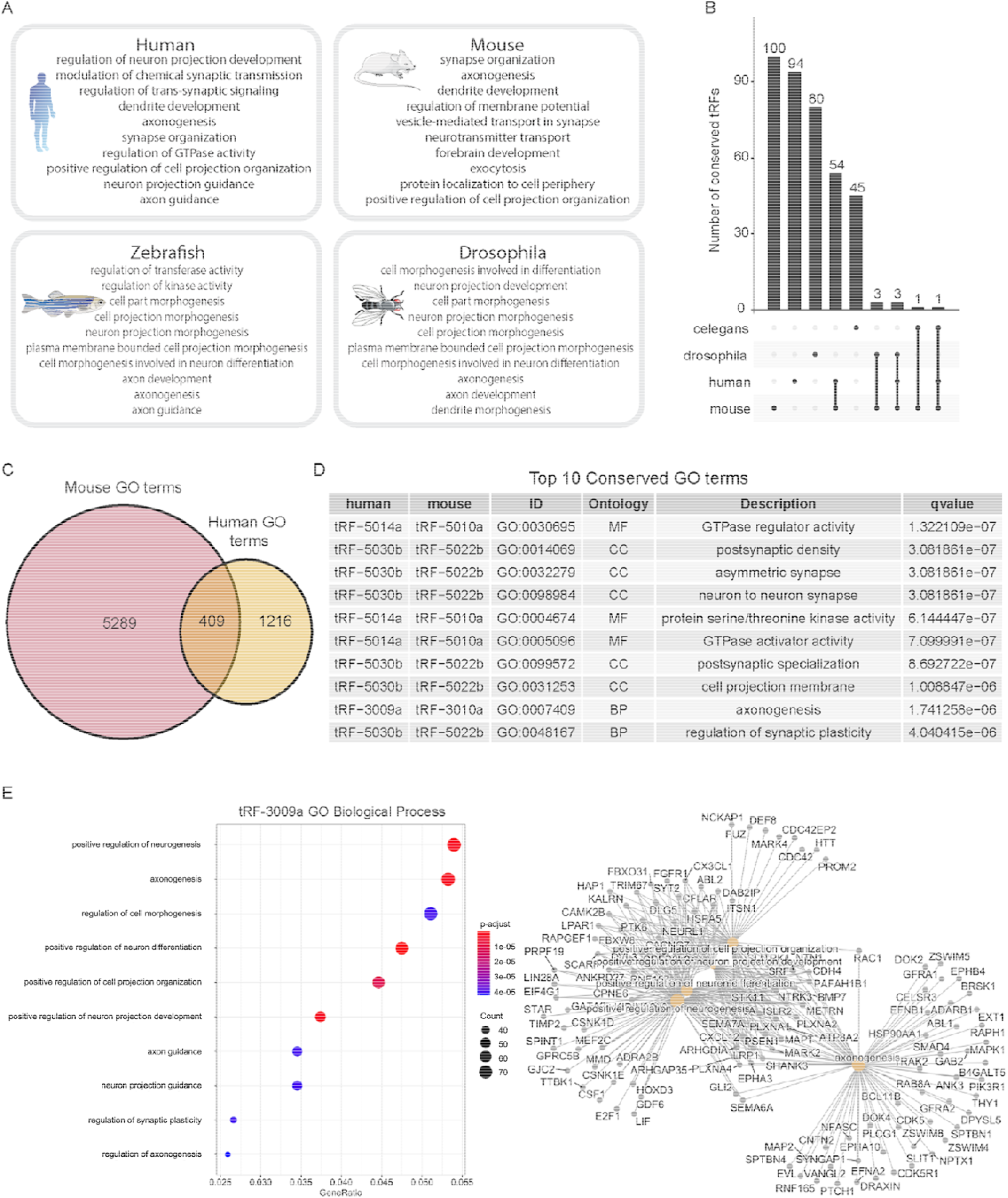
Several tRFs and tRF target GO terms are conserved. (A) Infographic showing the top 10 GO Biological Processes targeted by tRFs terms across four species. (B) Upset plot showing the intersection of conserved tRF sequences across four species. (C) Venn diagram of the intersection of conserved, significant GO terms of targets predicted for the 54 conserved tRFs between humans and mice. (D) Top ten GO terms conserved among predicted targets of human and mouse tRFs. Top ten GO terms were selected based on q value. MF: Molecular Function. CC: Cellular Component. BP: Biological Process. (E) An example gene ontology analysis plot describing biological processes enriched among predicted targets of tRF-3009a. Left: dot plot. Right: Gene-concept network plot.

tRForest users have the option to view and download the extended version of this plot with all three ontologies for any tRF and any species when possible. The dotplot allows users to quickly get a feel for the most probable pathways as well as the types of pathways affected for that tRF; for example, for human tRF-3009a it is evident from Figure 9 that the tRF may largely affect pathways related to neurons and their development. The plot contains information regarding gene ratio as the x-axis, adjusted p-value as dot color, and gene count as dot size, so the user can easily see the most relevant pathways as well as the underlying data, including statistical significance. The gene-concept network plot gives a more in-depth view of the connections between genes and pathways, which shows users not only how many genes influence a certain pathway, but also the level of connection and gene overlap between various pathways, which is not information given by the dotplot. Both of these plots give users an intuitive overview of their particular tRF, which can be very useful in looking beyond the genetic targeting capabilities of tRForest to how they can be applied. This also saves users the step of needing to perform GO themselves to see if a tRF is likely related to a certain biological process, molecular function, or cellular component.

### Database overview

Figure 2 provides a comprehensive workflow for the construction of the database. The database for tRForest contains predicted targets for seven species: human, mouse, *D. melanogaster, C. elegans, S. pombe, X. tropicalis*, and zebrafish. Notably, *R. sphaeroides*, which was included in tRFdb, is not included in tRForest because gene repression in this bacteria species is not Ago-mediated, as it is in the other species. In total, 628 tRFs with 262,880 target transcripts are included in the database. A more detailed breakdown by species is available on the Statistics page of the website. Predicted targets can be retrieved by filtering by tRF type or tRF ID as assigned in tRFdb; in addition, the database can be queried by gene name or Ensembl transcript ID to find tRFs that are involved in its repression. The output provided contains the tRF-ID, Ensembl transcript ID, gene name, binding energy of the tRF-gene duplex, binding location of the tRF on the 3’ UTR of the gene, an interaction illustration of the duplex, and is ordered by the prediction probability score given by tRForest which is also shown. The database also allows the user to download the 3’ UTR chromosomal coordinates of the predicted targets, as well as the output as a CSV file or Excel spreadsheet; GO analysis can also be viewed or downloaded. Further information about the tool, as well as statistics on the tool, a manual on usage, and a page to provide feedback can be found on the database’s webpage (https://trforest.com).

## Discussion

tRFs are becoming increasingly relevant in the study of biology and disease due to their involvement in regulating gene expression by pathways similar to miRNAs. However, there are few tools available that are specifically designed for tRF target prediction, and even fewer that utilize a machine learning-based approach. This study presents a comprehensive, rigorously designed and validated tRF target prediction tool in a convenient online database that includes tRFs across seven different species and includes tRF-1s, which other tRF algorithms neglect.

tRF targets were predicted using existing research on features involved in miRNA target prediction using random forests applied to tRF data. At the time of this study, there are several tRF targeting algorithms available, including tRFTar, tRFTarget, and tRFTars. However, only tRFTars uses a machine-learning approach, and tRForest is the first to use random forests in its algorithm. A significant advantage of using random forests is that they avoid overfitting, a common limitation of machine learning algorithms in which they become tailored specifically to the dataset they were trained on and thus become less predictive in independent datasets (24). This is evident by the greater effect size for tRForest predicted targets compared to tRFTars predicted targets in the independent validation datasets. In addition, it is the only tRF target prediction algorithm to generate corresponding negative sites for positive sites to better distinguish true targets, and the only one to return tRF-1 targets. Unlike several of the other algorithms, it also uses evolutionary conservation as a feature and provides a streamlined approach to accessing gene ontology information. In addition, it provides high-quality visualization of the enriched pathway information from the targets. As a result of this unique approach, tRForest matches or outperforms other existing tRF and miRNA target prediction tools for tRF target prediction.

tRForest has several limitations associated with data availability and efficiency. Feature calculation is currently rather time-consuming, so tRF targets are currently only available for existing tRFs found in tRFdb in species with Ago-mediated repression. In addition, there is a lack of available RNA-sequencing data with tRF overexpression to further validate the results. We attempted to address this limitation by testing tRForest on publicly available data, as well as data generated in our lab. The main limitation for the GO plot analysis is the fact that it requires a certain number of genes to generate plots (usually > 5), and for many tRFs this was not achieved with tRForest predictions. It was less of an issue for more popular model organisms, but S. pombe, for example, lacks many plots. This reduces convenience to users but is overall not a significant issue for the tool as a whole.

Improvements to the algorithm could be made by increasing the availability of CLASH and RNA-sequencing data. In addition, its efficiency can be increased by fixing the bottleneck in feature calculation which would allow the addition of a custom option to the tool, in which the user can input a unique tRF sequence and receive its targets.

This study presents a novel random forests-based machine learning model to predict tRF transcript targets in seven species. Furthermore, it provides a gene ontology analysis with these targets to determine enriched pathways. tRForest allows researchers to determine tRF targets and generate hypotheses about the biological functions of tRFs. The tool is available publicly at https://trforest.com.

## DATA AVAILABILITY

Sequencing data is available via GEO accession GSE189510.

## ACKNOWLEDGEMENTS

U87 cell was a kind gift from Roger Abounader (University of Virginia). We would like to thank University of Virginia Research Computing Core for computational. Icons in Figure 9A are from bioicons.com (patient icon by Marcel Tisch https://twitter.com/MarcelTisch is licensed under CC0; mouse-small icon by Servier https://smart.servier.com/ is licensed under CC-BY 3.0 Unported; Xenopus_laevis icon by DBCLS https://togotv.dbcls.jp/en/pics.html is licensed under CC-BY 4.0 Unported; drosophila-redeyes icon by Servier https://smart.servier.com/ is licensed under CC-BY 3.0 Unported).

## AUTHOR CONTRIBUTIONS

Conceptualization, B.W., R.P., F.H., and A.D.; Methodology, Investigation, Data Curation and Visualization, R.P., B.W., L.M., and Z.S.; Writing – Original Draft, R.P., B.W., L.M., Z.S., and A.D.; Writing – Review & Editing, B.W., Z.S., A.D.; Funding Acquisition, A.D., R.P., and B.W.; Supervision, A.D., F.H., and P.K.

