## Supplementary figures for "tRForest: a novel random forest-based algorithm for tRNA-derived fragment target prediction"

Supplemental figure 1 - Frequency of each nucleotide at each position in tRFs

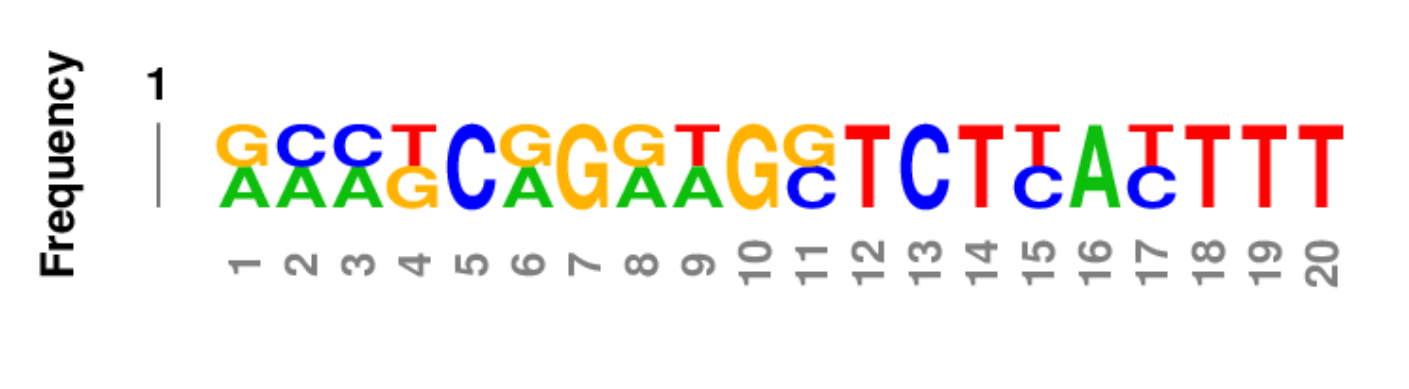

Supplemental figure 2 - CDF plots for tRF-3009a tRNA OE data in 293T (for five algorithms’ targets)

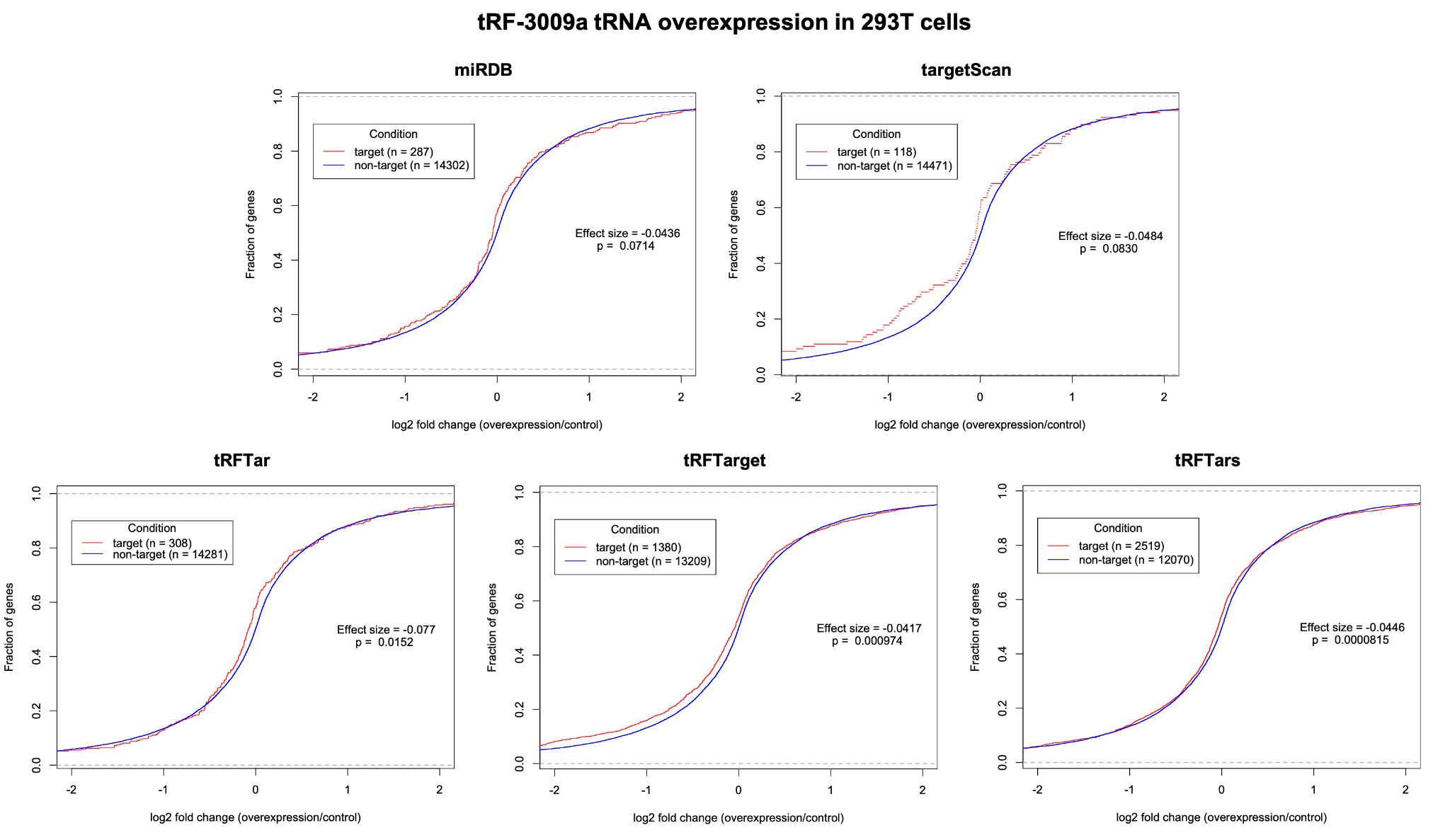

Supplemental figure 3 - CDF plots for tRF-3009a mimic OE data in u87 (for five algorithms’ targets)

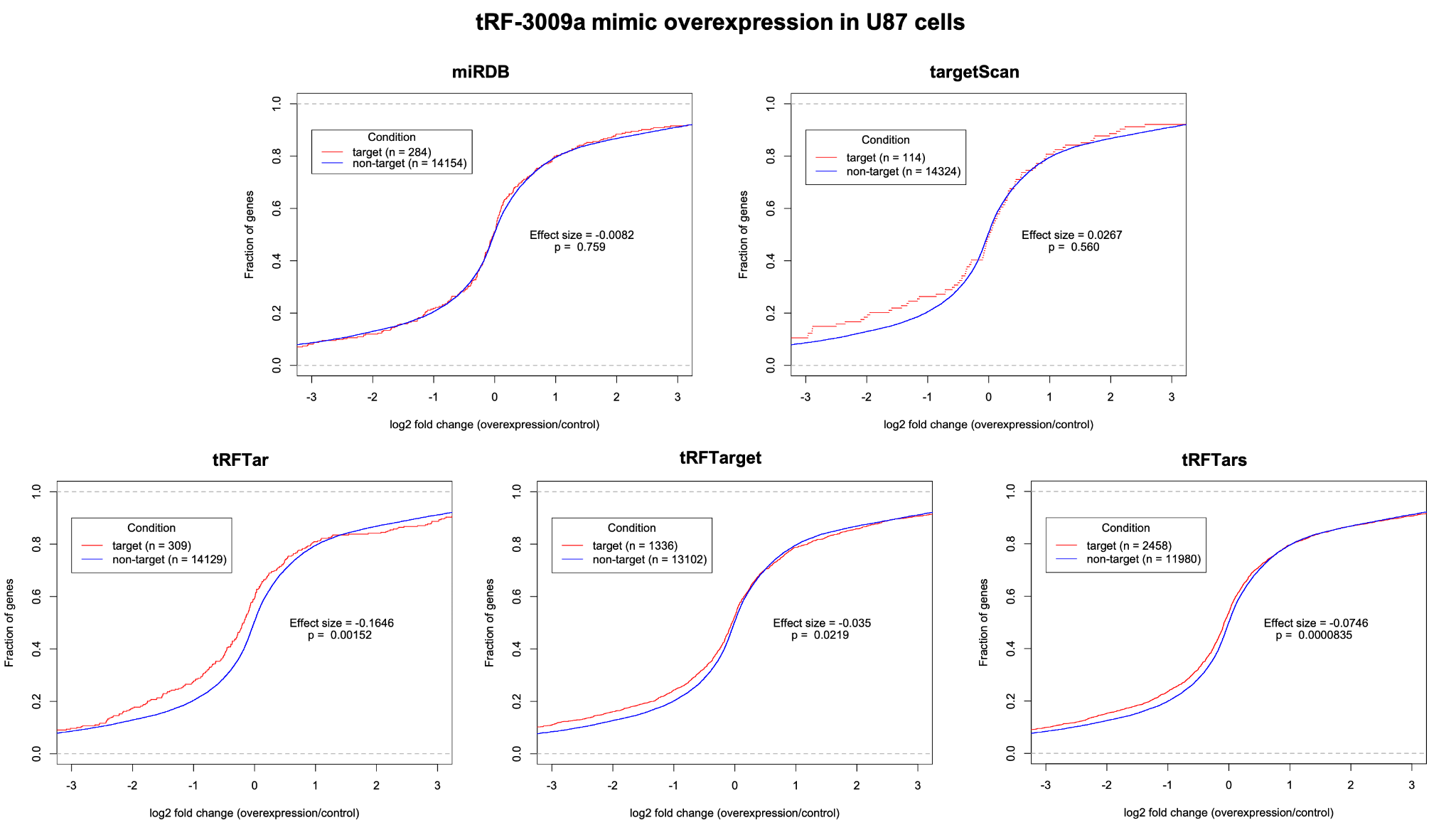
