## Supplementary file 1 for "tRForest: a novel random forest-based algorithm for tRNA-derived fragment target prediction"

**Feature calculation**

The following describes the method for calculating the features used by tRForest.

(1) **Binding energy**

This feature uses the RNAduplex function from the ViennaRNA package found at <https://www.tbi.univie.ac.at/RNA/>.

(2) **Seed match**

This is a binary feature (0 or 1) calculated by whether there existed perfect 7mer-m1 seed pairing (nucleotides 1-7 of the tRF).

(3) **AU content**

The local AU content around the binding site of the mRNA, 30nt upstream and downstream, given as a weighted sum by the proximity of the A/U to the binding site. The formula and a sample calculation for one direction is given below:

$$\sum_{i=1}^{30} \frac{P(i)}{i} =\frac{1}{1}+ \frac{0}{2}+\frac{0}{3}+\frac{1}{4}+\ldots+\frac{1}{29}=2.27546$$

P(i) is 1 if there is an A/U and 0 otherwise.

(4) **Number of paired positions**

This is just the count of the Watson-Crick pairings in the tRF-mRNA binding site.

(5) **Binding region length**

This is the length of the binding site in nucleotides.

(6) **Length of longest consecutive pairing**

This is the length of the longest consecutive Watson-Crick pairing region in the binding site.

(7) **Position of longest consecutive pairing**

This is the relative position of feature (6) to the 5’ end of the tRF.

(8) **Number of 3’ end pairs**

This is the count of Watson-Crick pairings in the last 7nt of the tRF (the 3’ end) binding site.

(9) **Difference between the number of paired positions in the seed and the 3’ UTR**

This is the difference between the number of Watson-Crick pairings in the seed (7 if the value of the seed match feature is 1) and feature (8).

**(10) Binding region conservation with phyloP**

This is the average phyloP score in the tRF-mRNA binding site. In this study, perBase phyloP scores were retrieved from UCSC. The required code used to obtain the scores is available on GitHub.

**(11) Flanking region conservation with phyloP**

This is the average phyloP score in 40nt upstream and downstream from tRF-mRNA binding site.

**(12) Binding region conservation with phastCons**

This is the average phastCons score in the tRF-mRNA binding site. In this study, phastCons scores were retrieved from UCSC.

**(13) Flanking region conservation with phastCons**

This is the average phastCons score in 40nt upstream and downstream from tRF-mRNA binding site.
